# A putative GtrB-like glycosyltransferase modulates cation-dependent BCP8-2 phage infection and cell surface structure in *Bacillus cereus*

**DOI:** 10.64898/2026.01.14.699495

**Authors:** Paul Tetteh Asare, Gaddapara Manasa, Yong Hwi Kwon, Seung Hyeon Ji, Myeong Seong Cha, Arun K. Bhunia, Jochen Klumpp, Kwang-Pyo Kim

**Affiliations:** Department of Food Science and Technology, College of Agriculture and Life Sciences, Jeonbuk National University, Jeonju 54896, Korea; Department of Agricultural Convergence Technology, College of Agriculture and Life Sciences, Jeonbuk National University, Jeonju 54896, Korea; Molecular Food Microbiology Laboratory, Department of Food Science, College of Agriculture, Purdue University, West Lafayette, IN 47907, USA; Institute of Food, Nutrition and Health, ETH Zurich, Schmelzbergstrasse 7, 8092, Zurich, Switzerland; Department of Food Tech, College of Agriculture and Life Sciences, Jeonbuk National University, Jeonju 54896, Korea; Connecticut State Department of Public Health, Rocky Hill, Connecticut, United States of America

**Author notes:** Corresponding Author: Kwang-Pyo Kim Department of Food Science and Technology, College of Agriculture and Life Sciences, Jeonbuk National University, Jeonju, Jeollabuk-do 54896, Korea. These authors contributed equally (Paul Tetteh Asare, Gaddapara Manasa).

## Abstract

*Bacillus cereus* is a foodborne pathogen of growing concern due to its persistence and antimicrobial resistance. To identify host determinants influencing susceptibility to phage BCP8-2, a mini-Tn*10* transposon mutant library of *B. cereus* ATCC 14579 was constructed and screened for altered phage susceptibility. A mutant (BC2012) showing partial resistance carried an insertion in *BC_RS27090* (previously *BC_5432*), encoding a putative GtrB-like bactoprenol glycosyltransferase. Complementation restored phage sensitivity, confirming its functional involvement. Adsorption and efficiency-of-plating assays revealed significantly reduced phage binding and infectivity in the mutant, particularly under cation-rich conditions. Microscopy showed altered surface morphology with thinner, smoother cell walls. These data indicate that *gtrB* affects surface properties essential for cation-dependent phage adsorption and infection and provide a foundation for future studies on the role of surface glycosylation and ionic interactions in Gram-positive phage biology.

**Importance:** Pathogenic *Bacillus cereus* is resilient across diverse environments and produces diverse toxins linked to foodborne outbreaks. Bacteriophages provide an effective strategy to control *B. cereus*; however, molecular targets of phage-host interaction are poorly understood, limiting the effective phage-based control. Current study addresses this gap by identifying a putative bactoprenol glycosyltransferase (GtrB), a vital factor in phage BCP8-2 susceptibility. Data obtained highlights partial reduction of phage infection in the *gtrB* mutant, demonstrating that precise glycosylation is essential for effective phage binding and entry. Our findings provide significant insights for advancing phage therapy, antibiotic resistance, and *B. cereus* biocontrol in food and clinical settings.

## Introduction

Phage adsorption, the initial attachment of viral particles to specific bacterial receptors, is a critical determinant of host specificity and successful infection (1). In diderm (Gram-negative) bacteria, adsorption often involves lipopolysaccharides (LPS), capsular polysaccharides (CPS), outer-membrane proteins (e.g., OmpC, OmpA, OmpF, LamB, PhoE, BtuB, FhuA), pili, and flagella (2–12). In monoderm (Gram-positive) bacteria, they typically rely on cell wall polysaccharides, including peptidoglycan-linked sugars (glucose, galactose, rhamnose, N-acetylglucosamine, N-acetylgalactosamine), wall teichoic acids (WTAs), lipoteichoic acids (LTAs), teichuronic acids, and membrane proteins such as PIP and YueB (13–18). Cell appendages (flagella, type IV pili) and surface-anchored proteins (e.g., GamR, CsaB) can also influence phage adsorption (19–21), illustrating the diversity of host determinants.

*Bacillus cereus* is a spore-forming foodborne pathogen capable of surviving harsh environmental conditions, including heat and desiccation, and causes diarrheal and emetic syndromes, whereas its relative *B. anthracis* can be fatal (15,16). Phage BCP8-2, a lytic Myovirus isolated from Korean fermented food, targets multiple *B. cereus* group species, including *B. cereus*, *B. mycoides*, *B. thuringiensis*, *B. weihenstephanensis*, and even *B. anthracis* (25, unpublished data). It lyses over 90% of food and clinical *B. cereus* isolates but does not infect beneficial species such as *B. subtilis* and *B. licheniformis* (26). Divalent cations enhance its infectivity, a feature observed in other phage systems (27). Its broad host range and specificity make BCP8-2 an ideal model for studying host determinants of phage susceptibility.

Here, we constructed a mini-Tn*10* transposon mutant library in *B. cereus* ATCC14579 to identify genes required for BCP8-2 infection. Screening revealed that disruption of *BC_RS27090* (previously *BC_5432*; designated as *gtrB* hereafter), encoding a putative GtrB-like bactoprenol glycosyltransferase, significantly reduced phage adsorption and infectivity. Complementation restored susceptibility, confirming the gene’s role in phage–host interactions. Microscopic analyses, including Gram staining, transmission electron microscopy, and field-emission scanning electron microscopy, revealed altered cell wall morphology and increased autolysis. These findings highlight *gtrB* as a key determinant of cell wall integrity and efficient phage adsorption in *B. cereus*, providing new insights into phage–host interactions and potential biocontrol applications.

## Materials and methods

### Bacterial strains, phages, plasmids, primers and bacterial growth conditions

All bacterial strains, phage, and plasmids used in this study are listed in **Table 1** and primers are provided in **Table 2**. *Escherichia coli* and *B. cereus* ATCC 14579 were grown in Luria Bertani media (LB; Oxoid, Detroit, MI) and tryptic soy broth (TSB; Difco), respectively, at 37°C, unless otherwise indicated. When needed, antibiotics were added at 4 µg/mL of erythromycin and 100 µg/mL of spectinomycin. Phage BCP8-2 was propagated in *B. cereus* JCM2152 as described previously (22).

**Table 1.**
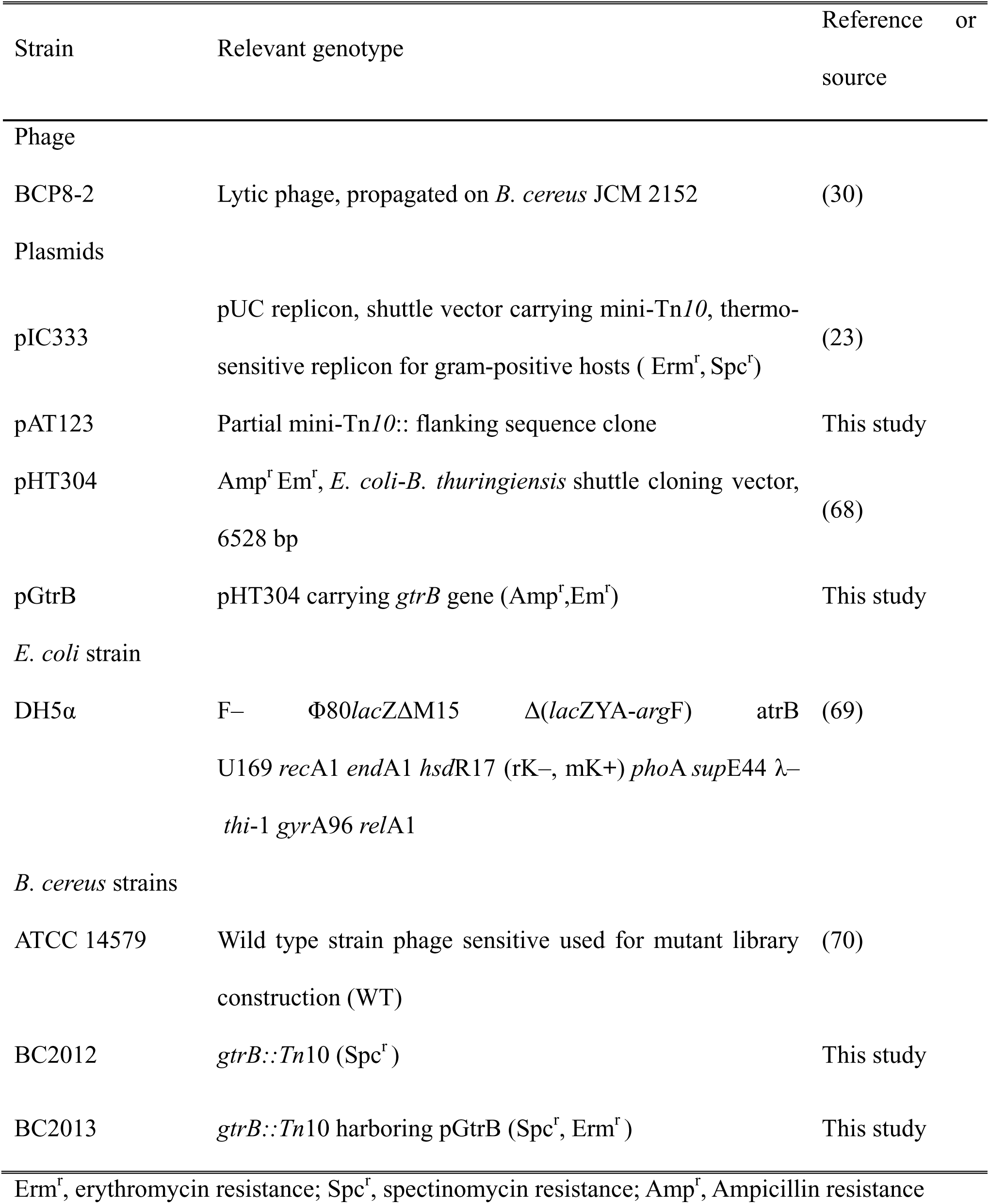
Phages, plasmids and bacterial strains used.

**Table 2.**
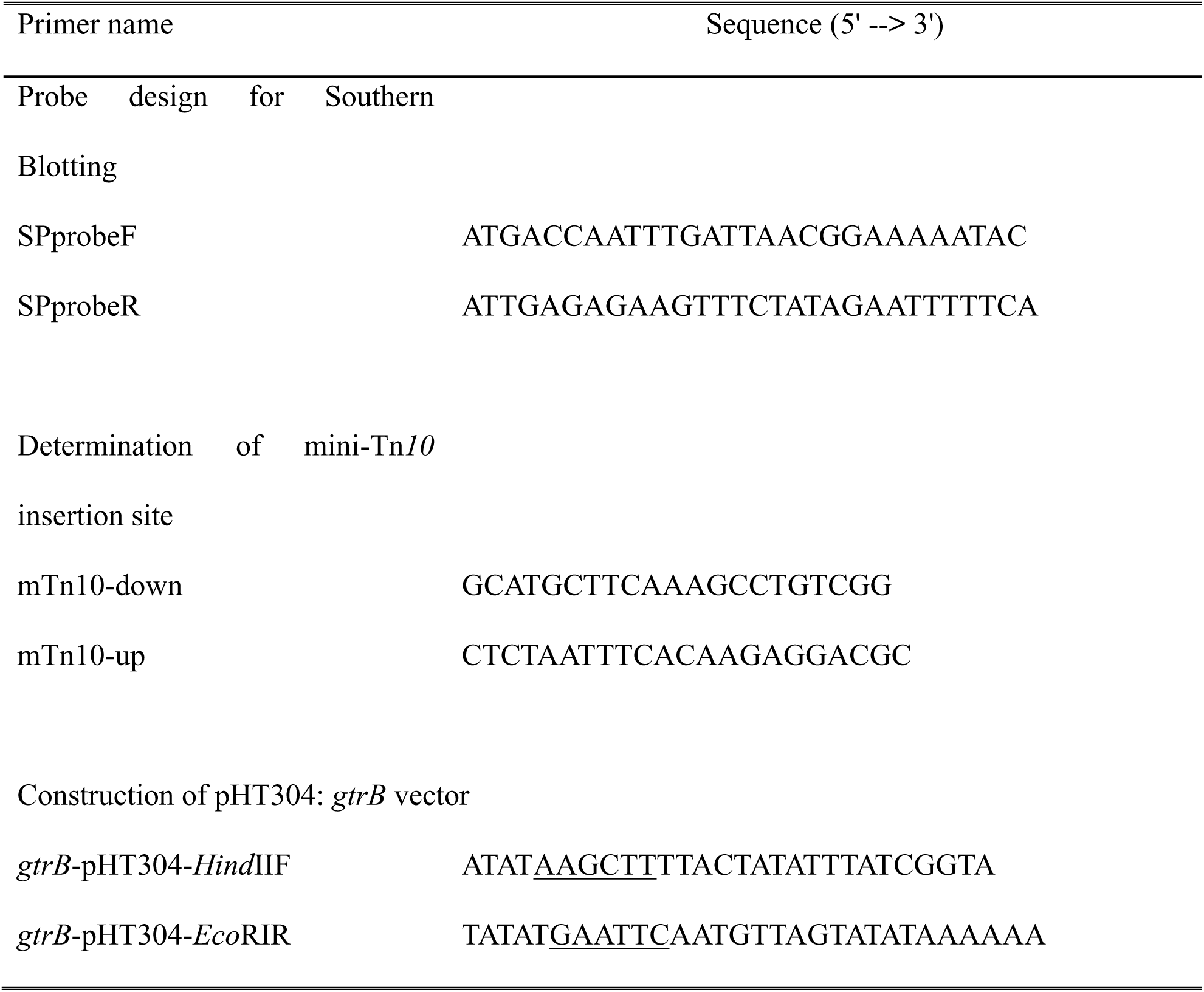
Primers used for PCR amplification and sequencing.

### Transposon mutagenesis

A transposon mutant library in *B. cereus* ATCC 14579 was constructed using the mini-Tn*10* transposon in the pIC333 plasmid (23). Preparation of electro-competent cells and transformation with pIC333 were performed as described previously by Turgeon et al., 2006 (24) and Steinmetz et al., (1994) (25). Transformants were selected for Spc^R^ and Erm^R^ at permissive temperature (30°C). To induce transposition, the isolated clones were subjected to plasmid curing as described by Wilson and Szurmant, 2011 (26) and Spc^R^ and Erm^R^ mutants were selected to prepare the library.

### Southern hybridization

Genomic and plasmid DNA isolations were performed using the Exgene Cell SV mini kit (GeneAll, Seoul, Korea) and the QIAGEN Plasmid Mini Kit (Qiagen, Valencia, CA), respectively. Southern blot analysis was carried out as described by Veeranagouda et al., 2012 (27). Genomic DNA (2 µg), digested with *Eco*RI was separated by agarose gel electrophoresis (0.7% agar) and transferred to positively charged nylon membrane (Pall Corporation, Port Washington, NY) using the downward capillary transfer system as described by Sambrook et al., 1989 (28). DNA was cross-linked to the membrane by baking at 80°C for 2 h. The probe specific for the spectinomycin-resistance gene was PCR-amplified using SPprobeF and SPprobeR primers, and pIC333 as the template DNA. The amplified DNA was labelled using the Random Primed DNA Labeling system (Roche Diagnostics, Basel, Switzerland) and hybridization and detection of the probe were performed with the DIG DNA Labeling and Detection Kit (Roche Diagnostics) as described by the manufacturer.

### Screening of phage-resistant mutants

We employed the concept of reverse phage typing as described previously by Loessner et al., (29) to screen phage-resistant mutant(s). Briefly, a mixture of 100 µL of phage stock (10^7^ PFU [plaque forming unit] /mL) and 4 mL of TA soft agar (0.4% agar in TA broth; 8 g nutrient broth, 5 g NaCl [86 mM], 0.2 g MgSO_4_·7H_2_O [0.8 mM], 0.05 g MnSO_4_ [0.3 mM], and 0.15 g CaCl_2_ [1.0 mM] per 1 L, pH 5.9–6.0) (30). was poured on LB agar plates (1.2% agar) and allowed to solidify. Thirty minutes later, 5 µL of overnight-grown bacterial culture was spotted on the plates. The plates were dried for 10 min at room temperature and then incubated at 37°C for 12 h. Mutants (including BC2012) that formed a bacterial lawn in the presence of phage BCP8-2 were further characterized.

### Identification of transposon insertion site

The transposon insertion site was determined as previously described (29). Briefly, genomic DNA digested with *Eco*RI (does not cut the transposon insertion sequence) was self-ligated and introduced into *E. coli* DH5α by electroporation. Spectinomycin-resistant *E. coli* were recovered and then the plasmid was sequenced, using transposon-specific outward primers mTn*10*-down and mTn*10*-up (**Table 2**). The sequence was then compared to the nucleotide database (GenBank accession number NC_004722) of *B. cereus* ATCC 14579.

### Bioinformatics

The Basic Local Alignment Search Tool (BLAST) (31) and Conserved Domain Database (CDD) programs were employed to perform a protein BLAST and conserved domain (32) search, respectively, through the public interface at (http://blast.ncbi.nlm.nih.gov/Blast.cgi) using default settings (version 2.2.17). Transmembrane segments and membrane topology were predicted using TMHMM (http://www.cbs.dtu.dk/services). Operons were predicted using the MicrobeOnline database (http://microbesonline.org) (33).

### Complementation study

Complete open reading frame (939 bp) and 300 bp upstream to putative bactoprenol glycosyl transferase (*gtrB*) gene in *B. cereus* ATCC 14579 were amplified using *gtrB*-pHT304-*Hind*IIIF and *gtrB*-pHT304-*Eco*RIR primers (**Table 2**) with the PCR conditions of holding at 95°C for 5 min, followed by amplification for 30 cycles at 95°C for 60 s, 50°C for 60 s, and 72°C for 3 min, with a final extension for 10 min. The PCR fragment was then digested with *Hind*III and *Eco*RI, cloned into pHT304 vector (34) (which was also digested with both enzymes) and transformed into the mutant BC2012 to construct BC2013 (the complement strain).

### Phage dotting assay

The mixture of 200 µL of overnight bacterial culture and 4 mL of TA soft agar (0.4%) was poured onto LB plates to prepare a bacterial lawn, which was then allowed to solidify for 30 min. Later, 10 µL of phages diluted in SM buffer (50 mM Tris–HCl, pH 7.5, 100 mM NaCl, 10 mM MgSO_4_) (30) were dotted on the lawn, allowed to dry at room temperature for 10 min. The plates were then incubated at 37°C for 10 h, after which the plaques or lysis zones were inspected. Plate images were recorded using CoreImager (CoreBio, La Jolla, CA).

### Bacterial growth inhibition by phage in liquid culture

The bacterial lysis assay was performed as described previously (35) with modifications in media type. Nutrient broth (NB) or TA broth (25 mL) was inoculated with 1% overnight culture of *B. cereus* JCM2152, and incubated at 37°C until the culture reached OD_600_ (optical density at 600 nm) of 0.5 (equivalent to 10^7^ CFU/mL). Then 250 µL of phage stock was added at an MOI of 1 or 0.1. The mixture was incubated statically for 10 min at 37°C and incubated at 37°C with shaking (160 RPM). The OD_600_ was monitored every 2 h to monitor bacterial growth.

### Adsorption assay

The phage adsorption assay was performed as previously described by Dupont et al. (36) with modifications. Cells were grown in either NB or TA broth until OD_600_ reached 0.5. Then, cells were infected with phage at an MOI of 0.001 and incubated for 10 min at 37°C. After centrifugation (15000 x g, 4°C, 3 min), unadsorbed phages in the supernatant were counted by a standard plaque-forming assay, plating with the propagation host, *B. cereus* JCM2152. The percentage adsorption was calculated as follows: [1-(phage titer of the supernatant after centrifugation/phage titer of a control reaction supernatant after centrifugation] x 100.

### Efficiency of plating

Overnight-grown bacterial culture (200 µL) and serially diluted BCP8-2 (100 µL) were added to 4 mL of TA soft agar and poured onto pre-solidified LB agar plates, which were incubated at 37°C for 12 h, after which PFUs were counted (30).

### Gram staining

Staining of the strains were performed as described previously (37).

### Transmission electron microscopy (TEM)

Transmission electron microscopy (TEM) was performed as previously described (38) with few modifications. Overnight culture (1%) of *B. cereus* JCM2152 was used to inoculate fresh LB media (25 mL) and incubated at 37°C for 12 h. After centrifugation at 10,000 x g for 5 min, the pellet was fixed with 1 mL of a modified Karnovsky’s fixative (2% paraformaldehyde and 2% glutaraldehyde in 0.005 M sodium cacodylate buffer, pH 7.2) for 2 h at 4°C. The sample was washed three times with 0.05 M sodium cacodylate buffer (pH 7.2) for 10 min at 4°C and treated with 1% osmium tetroxide in 0.05 M sodium cacodylate buffer (pH 7.2) for 1.5 h at 4°C. The sample was washed twice for 5 min with distilled water at room temperature, followed by en-bloc staining 4°C overnight using 0.5% uranyl acetate. The sample was dehydrated through graded ethanol series (30 to 100%), and was embedded in a mixture of propylene oxide and Embed 812 resin. Finally, the sample was dried, sectioned and stained with uranyl acetate (0.5%). The sample was examined under Hitachi H-7650 TEM (Japan) at an acceleration voltage of 100 kV, and images were recorded using conventional electron imaging films.

### Field emission scanning electron microscopy (FE-SEM)

Sample preparation for FE-SEM analysis was performed as described for TEM (38) with some modifications. After the dehydration step, the samples were treated with 100% HMDS (hexamethyldisilazane) for 15 min and allowed to air dry overnight and observed using SIGMA VP Field Emission-SEM (Zeiss, Oberkochen, Germany) at an acceleration voltage of 100 kV.

### Autolysis study

Autolysis experiment was conducted as described by Falk et al. (39) with slight modifications. Briefly, overnight culture (1%) was used to inoculate fresh LB media (25 mL) and incubated at 37°C with (160 RPM) and without shaking (39). OD_600_ was measured every 2 h and CFU counting every 6 h to monitor bacterial growth.

## Results

### A putative bactoprenol glycosyltransferase (*gtrB*) homolog is disrupted in a BCP8-2 resistant mutant

To identify host factor(s) essential for BCP8-2 infection, a library of 4,200 random transposon mutants was constructed in *B. cereus* ATCC 14579 using pIC333 and screened for reduced phage susceptibility. Twenty-three mutants were randomly selected and validated by Southern hybridization using a spectinomycin resistance gene probe. All 23 contained a single transposon copy, and 18 distinct fragment sizes indicated predominantly random single-copy insertions (Supplementary figure 1).

Screening identified a mutant BC2012, which displayed impaired phage infection across multiple parameters (see below). The transposon insertion site was mapped via plasmid recovery to generate pAT123. Sequencing of the flanking genomic DNA followed by BLAST search revealed insertion within the *BC_RS27090* gene (939 bp) in *B. cereus* ATCC14579, 48 bp downstream of the start codon, and Southern hybridization confirmed the single specific insertion (data not shown).

Bioinformatic analysis annotated BC_RS27090 as a putative bactoprenol glycosyltransferase (GtrB) (40). Its deduced amino acid sequence shares 42.43% identity with *Shigella flexneri* GtrB (E-value 2e-84) (41). Conserved domain and BLASTp analyses identified a DMP1-like bac domain (cd04187; E-value 5.22e-90) at the N-terminus, which belongs to the Glyco_tranf_GT-A_type superfamily (cl11394), and contains a single DXD motif (cd04187), indicative of the sole active site and characteristic of metal-dependent GT-A glycosyltransferase (42). TMHMM predicted two C-terminal transmembrane domains, as observed in *Shigella flexneri* GtrB, suggesting a role in membrane association and access to lipid-linked sugar substrates (UDPs) (43). MicrobeOnline analysis indicated that BC_RS27090 functions as a single-gene unit.

### The *gtrB* gene product is required for BCP8-2 infection

Phage infection was assessed in WT strain, the transposon mutant (BC2012), and the complement strain (BC2013) in TA (cation-supplemented) and NB broths (Figure 1). In TA at MOI 1, WT cells lysed rapidly, with OD₆₀₀ decreasing from 0.5 to 0.02 within 6 h. At MOI 0.1, partial inhibition occurred (Figure 1A). In contrast, BC2012 showed no growth inhibition at either MOI (Figure 1B). Complementation of *gtrB* restored WT-like lysis in BC2013 (Figure 1C). In NB broth, lysis in WT was slower and less pronounced; at MOI 0.1, no significant inhibition occurred, and MOI 1 required over 6 h to reduce OD₆₀₀ below 0.1 (Figure 1D). BC2012 showed no significant growth inhibition by BCP8-2 (Figure 1E), while the complement strain BC2013 displayed identical growth patterns to the WT (Figure 1F)

**Figure 1.**
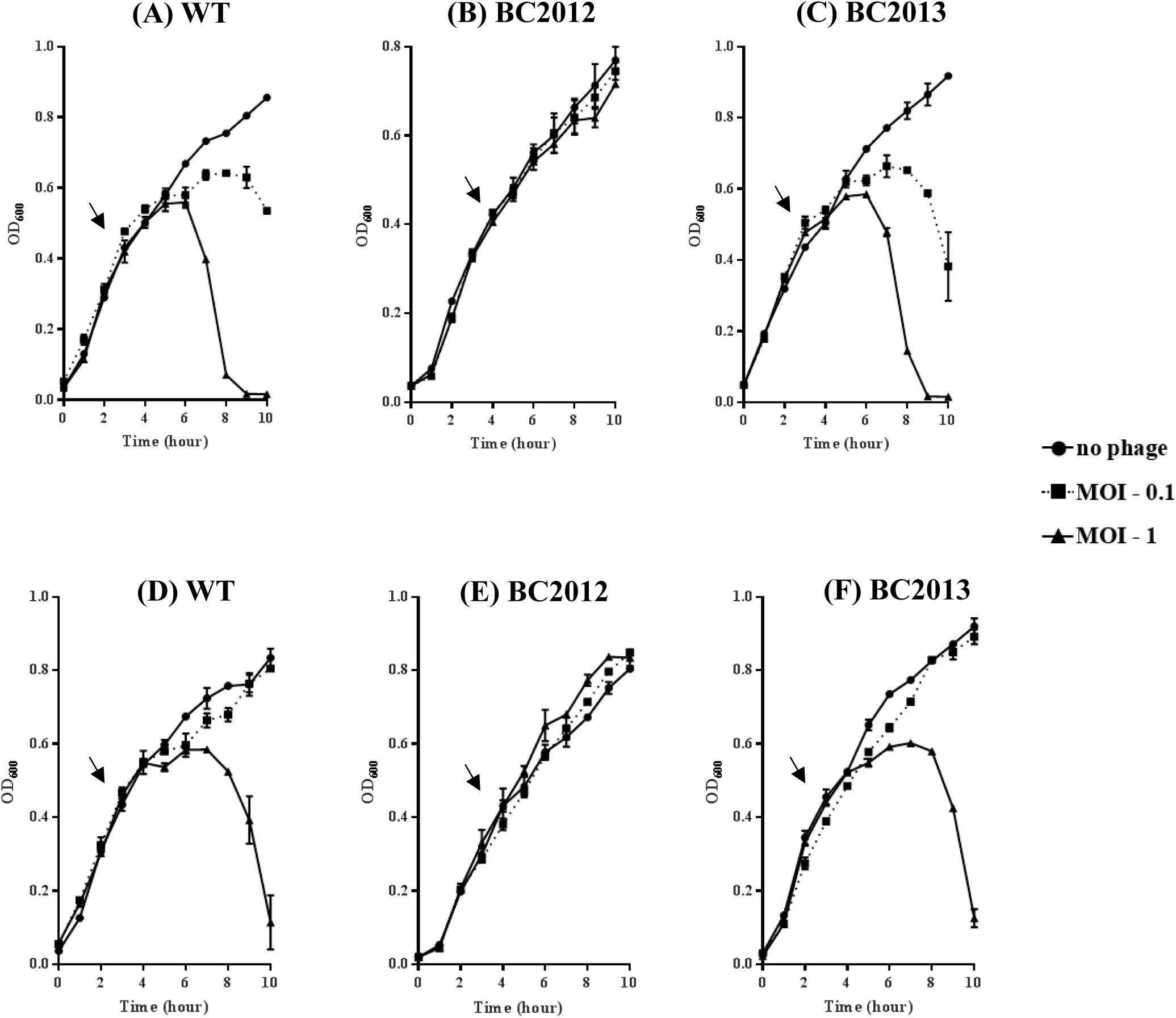
Inhibition of *B. cereus* strains in liquid culture by BCP8-2. wildtype (WT), the mutant BC2012, and the complementation strain BC2013 at MOI 1 and 0.1 with no phage as control in TA (1A, 1B, 1C) and NB (1D, 1E and 1F). The arrow indicates the point of phage addition.

Spot-on-lawn assays confirmed these findings: WT and BC2013 produced clear zones of inhibition at phage titers as low as 10⁵ PFU/mL, whereas BC2012 required ≥10⁷ PFU/mL to achieve similar inhibition (Supplementary Figure 2).

### *gtrB* mutation decreases BCP8-2 adsorption and efficiency of plating (EOP)

Adsorption assays at MOI 0.001 over 10 min revealed significant differences. In NB, WT adsorbed 60.4% of phages, BC2012 only 4.64%, and BC2013 partially restored adsorption (51.16%). In TA, adsorption increased to 94.84% in WT and 90.47% in BC2013, while BC2012 remained low (4.19%) (Figure 2), indicating the importance of divalent cations in *gtrB*-mediated phage adsorption.

**Figure 2.**
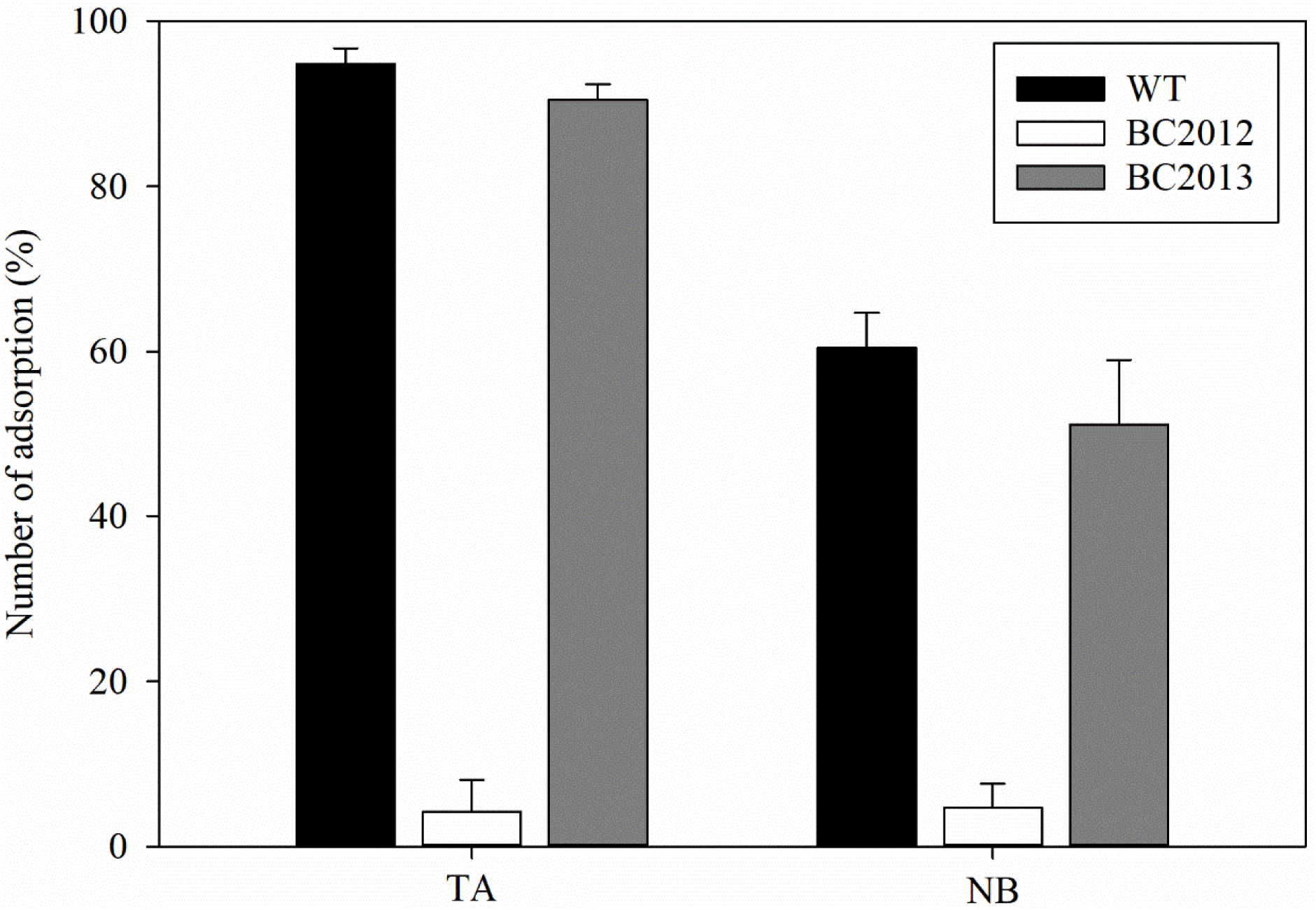
Adsorption assay of BCP8-2 reduced in the *gtrB* mutant. Adsorption rate of BCP8-2 to Wild-type (WT), the mutant BC2012, and the complement BC2013 using TA (Cation rich NB) and NB (Nutrient broth) media. Phage is added to cultures at MOI 0.001 when OD600 is 0.5. Relative phage titer in supernatant over time was determined using the spot assay.

EOP assays using soft agar overlays showed that WT in TA agar yielded 8.45 × 10⁹ PFU/mL (100%), BC2012 produced 2.85 × 10⁹ PFU/mL (∼34%), and the complement BC2013 restored EOP to 98% (8.30 × 10⁹ PFU/mL). In NB agar, EOP of WT and BC2013 was lower, and BC2012 matched WT, reinforcing the role of cations in plaque formation (Figure 3).

**Figure 3.**
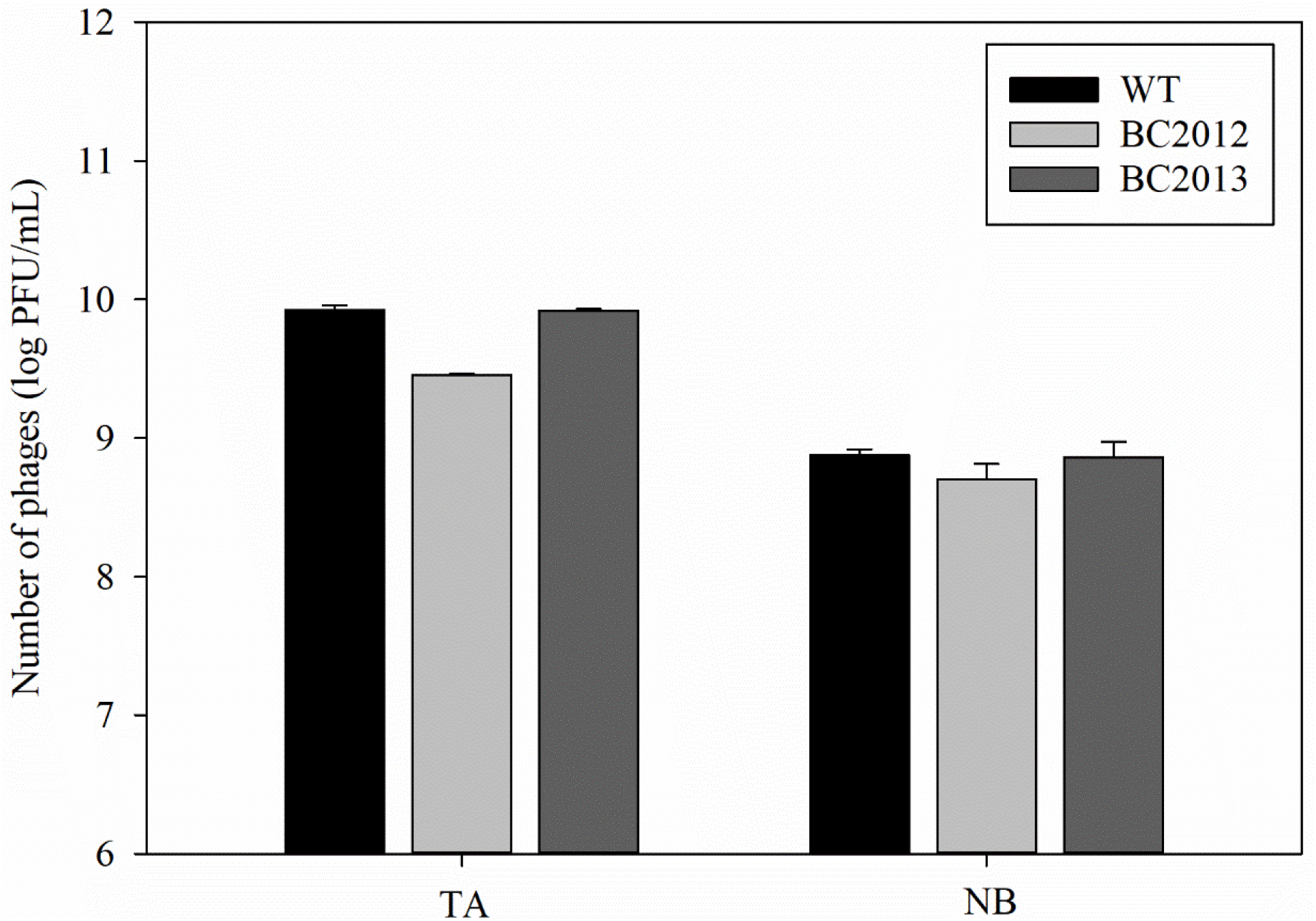
Efficiency of plating (EOP) of phage BCP8-2. Comparison of plating efficiency of phage BCP8-2 on wildtype (WT), the mutant BC2012, and the complement BC2013 strains using double agar-overlay assay in TA-based and NB-based media.

### *gtrB* mutation affects BCP8-2 plaque morphology in *B. cereus*

Plaque analysis revealed WT and the complement BC2013 produced smaller, rough-edged plaques, whereas BC2012 formed larger, smooth-edged plaques (Figure 4). The diffused plaque morphology is often associated with low adsorption rate (44), consistent with its reduced adsorption rates in BC2012.

**Figure 4.**
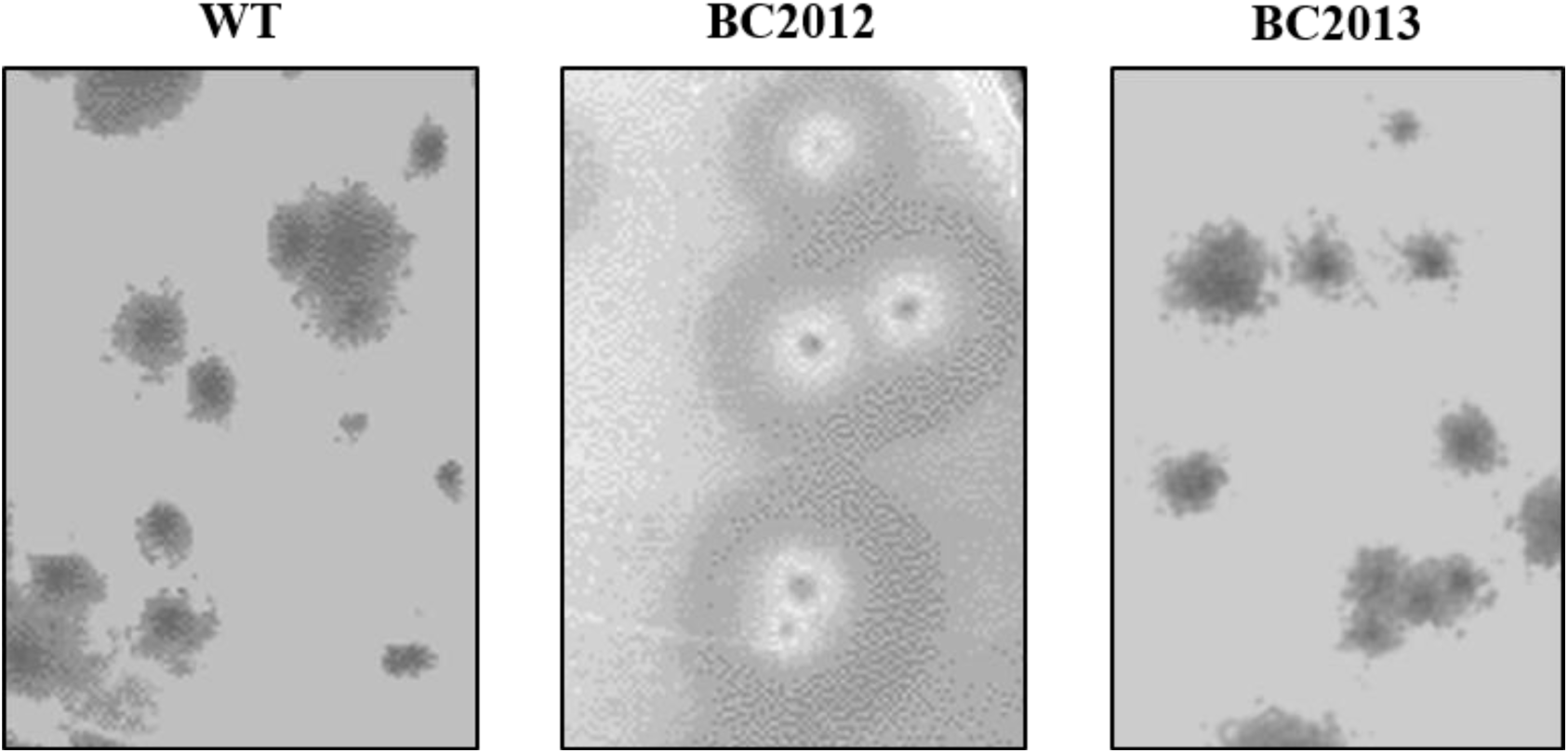
*gtrB* mutation affects BCP8-2 plaque morphology in *B. cereus*. Plaque morphology of phage BCP8-2 with wildtype (WT), the mutant BC2012, and the complement BC2013. The mutant strain BC2012 formed larger diffused plaques compared to the smaller and rougher WT and the complement BC2013 strains.

### *gtrB* mutation affects the bacterial surface structure of *B. cereus*

Morphological analyses using Gram staining, TEM, and FE-SEM highlighted substantial structural changes in BC2012 surface to WT and the complement BC2013 (Figure 5). Gram staining showed that BC2012 cells appeared pink, indicating a Gram-negative-like phenotype, whereas WT and BC2013 retained the typical purple coloration of Gram-positive bacteria (Figure 5A). TEM revealed that the BC2012 cell wall was significantly thinner than that of WT and the complement (Figure 5B), suggesting compromised peptidoglycan integrity.

**Figure 5.**
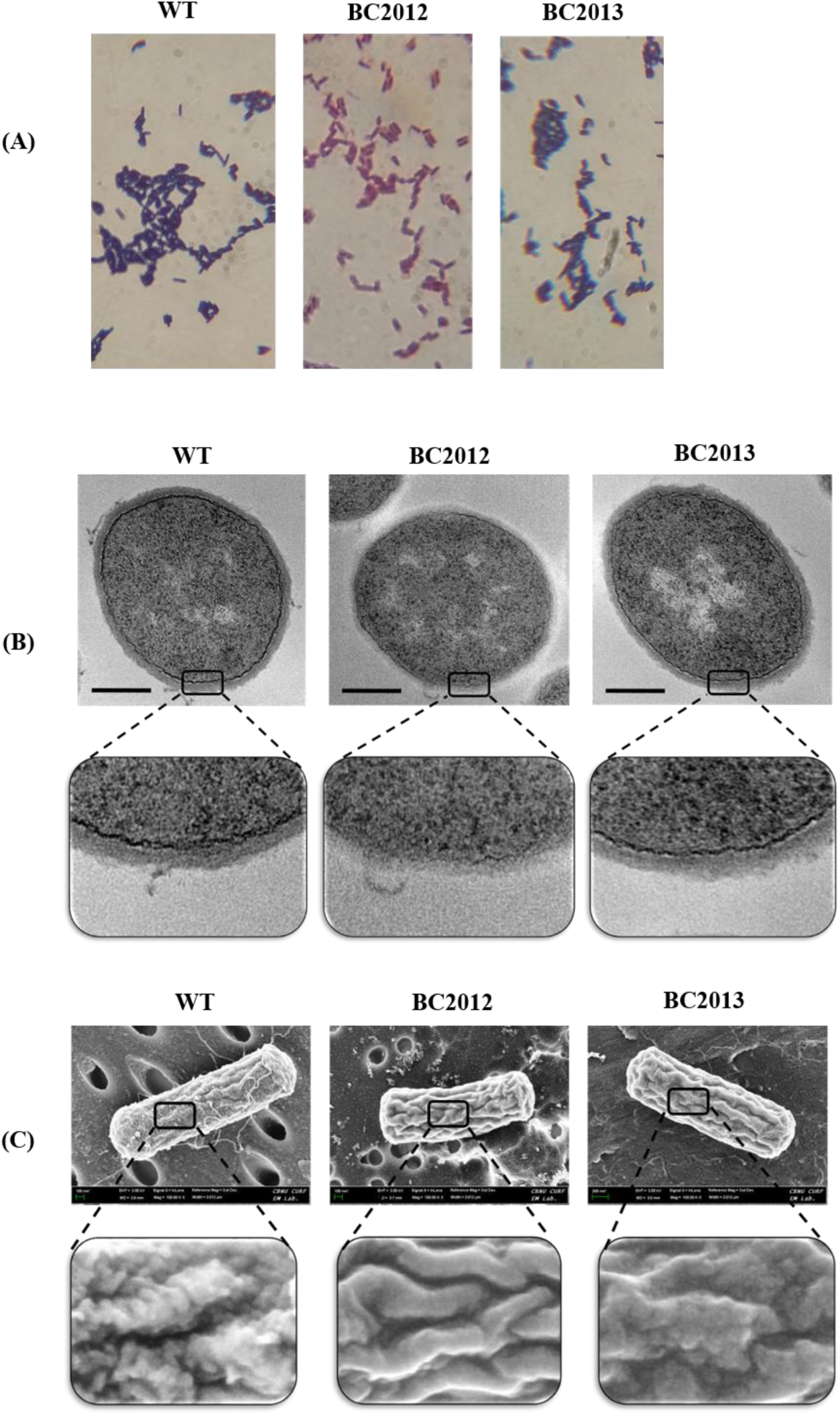
*gtrB* mutation affects the bacterial surface structure of *B. cereus*. Effect of *gtrB* mutation on (A) Gram staining, magnification ×1000 (B) Transmission electron micrographs of *B.cereus* (scale bar 200 nm) and (C) FE-SEM images of the WT, the mutant BC2012 and the complement BC2013 (at 100kV).

FE-SEM provided further insight into surface architecture: WT cells displayed long rods with bumpy, rough surfaces and numerous folds, while BC2012 cells were comparatively smooth with fewer surface folds, reflecting a potential loss of structural complexity (Figure 5C). Complementation partially restored rough surface morphology. Together, these findings indicate that *gtrB* is essential for proper assembly or modification of cell wall polysaccharides, which likely serve as key receptors for BCP8-2 adsorption. Alterations in surface structure may not only reduce phage-binding efficiency but also contribute to increased susceptibility to autolysis, as observed in static growth conditions (see below).

### Transposon insertion in *gtrB* induces autolysis

Time-lapse experiments under static conditions showed BC2012 cultures became clear after 24 h, whereas WT and BC2013 cultures remained turbid (Figure 6A). Under shaking (160 RPM), no differences were observed (Figure 6B). CFU counts over 48 h confirmed continuous decline in BC2012 under static conditions (1.41 × 10⁶ CFU/mL at 48 h) versus stable counts for WT (1.66 × 10⁸ CFU/mL) and BC2013 (1.26 × 10⁸ CFU/mL) (Figure 7A–B).

**Figure 6.**
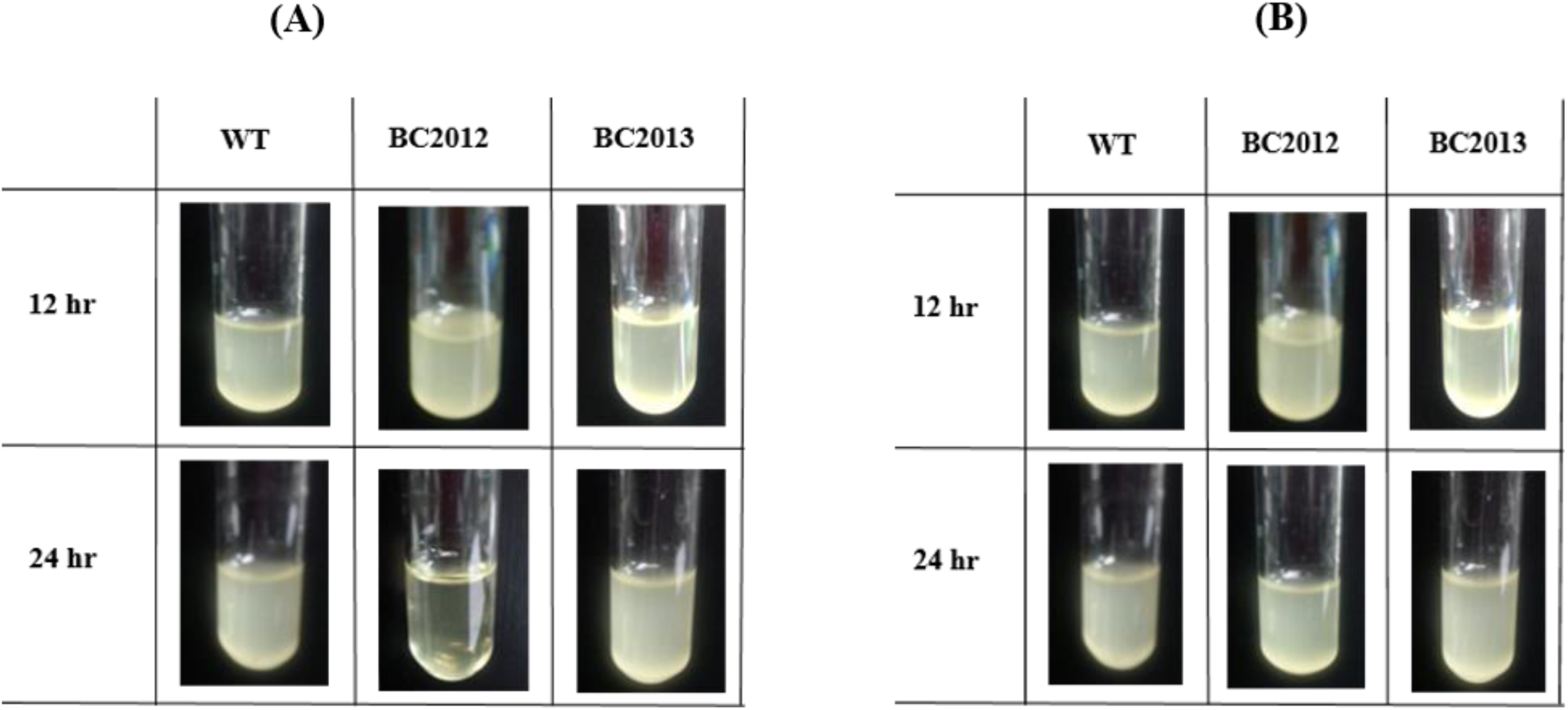
**Autolysis assay** of the WT, the mutant BC2012, and the complement BC2013 under (A) static and (B) shaking conditions, respectively. (160 rpm, 12 h and 24 h incubation)

**Figure 7.**
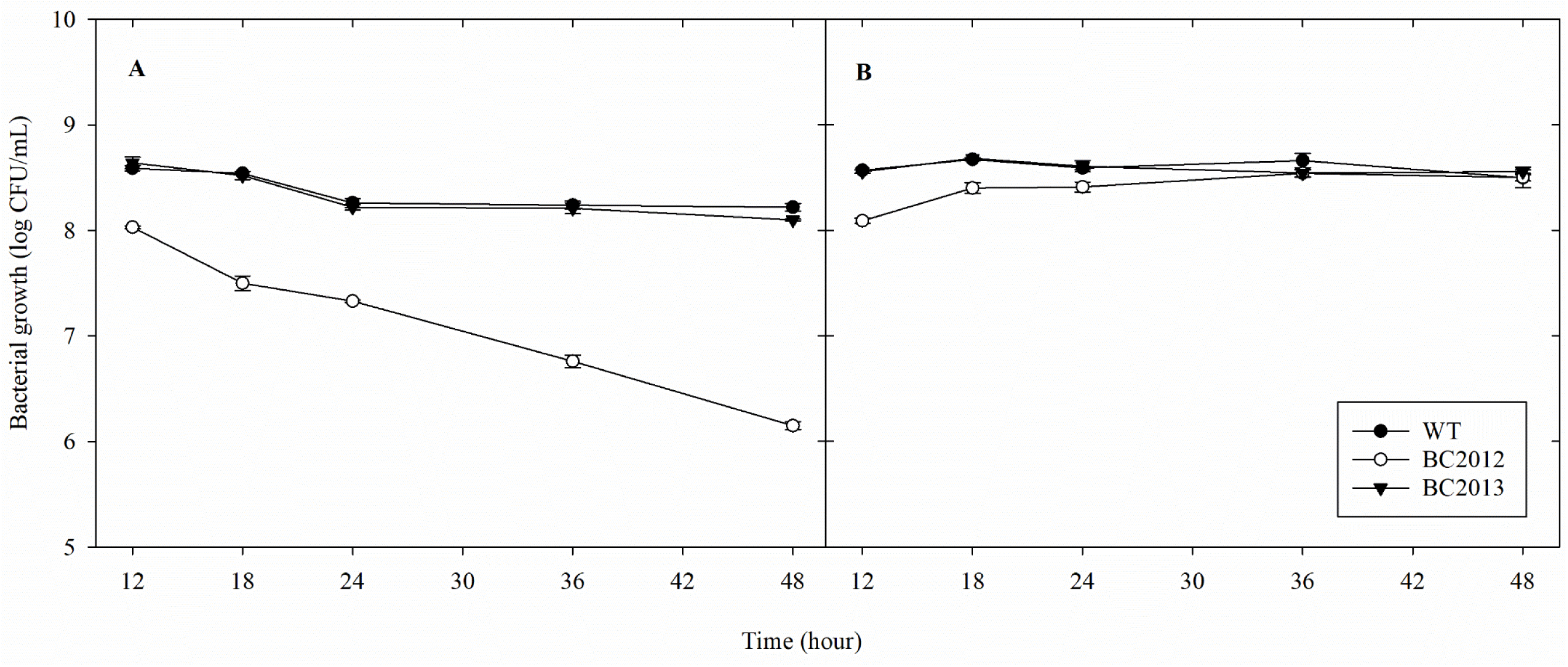
Transposon insertion in *gtrB* induces autolysis. Growth characteristics of the WT, the mutant BC2012, and the complement BC2013 under static (A) and shaking (B) conditions (160 rpm, 48 h incubation). Data shown were the mean of three independent experiments.

In addition, mutant colonies displayed central lysis zones, absent in WT and BC2013 (data not shown), consistent with possible autolysis.

## Discussion

In this study, we identified a *gtrB* homolog (*BC_RS27090*), encoding a putative bactoprenol glycosyltransferase, as an important determinant of *B. cereus* ATCC 14579 susceptibility to phage BCP8-2. Screening a random transposon mutant library revealed BC2012, carrying an insertion in *gtrB*, which exhibited marked resistance to BCP8-2 infection. Complementation restored wild-type sensitivity, confirming the functional role of *gtrB* (Fig. 1A–F). Glycosyltransferases like GtrB are integral to the synthesis of bacterial surface glycopolymers, including wall teichoic acids (WTAs) and lipoteichoic acids (LTAs), which serve as frequent targets for phage adsorption in Gram-positive bacteria (45–47). The identification of *gtrB* as a host factor establishes a link between cell envelope glycosylation and phage susceptibility in *Bacillus*, a genus where phage receptor mechanisms remain poorly characterized.

GtrB has been implicated in phage resistance in Gram-negative bacteria, notably *Shigella flexneri*, where GtrB-mediated O-antigen glucosylation modifies LPS structures that serve as phage receptors and enable evasion of phage recognition (41–48). To our knowledge, its role in Gram-positive bacteria, including *Bacillus* spp., has not been reported.

Previously, TagE, a glycosyltransferase in *B. subtilis*, was reported to participate in WTA glycosylation and influence phage infection (45,49,50). TagE contains a Glycosyltransferase_1 domain A (pfam09318; E-value 9.14e-117) with a GTB-type fold (cl10013) and transfers glucose from UDP-glucose to the α-carbon of poly-glycerol phosphate. In contrast, bioinformatic analyses of *B. cereus* GtrB homolog (312 amino acids) indicate a Glycosyltransferase 2-like domain (IPR001173) with a DXD motif that interacts with divalent cations (48,51). Proteins of this family catalyze metal-dependent sugar transfer from activated donors to lipid or carbohydrate acceptors, contributing to O-antigen, LPS, WTA, LTA, CPS, and glycan modifications (48,52–56), potentially influencing phage receptor structures (41). Thus, although both TagE and GtrB-homolog are glycosyltransferases, they belong to distinct structural folds and GT families, indicating different catalytic mechanisms (57).

Our results show that disruption of *gtrB* in *B. cereus* affects BCP8-2 phage adsorption, plating efficiency, and plaque morphology, indicating a functional role in the formation or modification of a surface receptor required for efficient phage attachment (Figs. 2–4). Phage adsorption assays revealed that BC2012 had dramatically reduced adsorption (<5%) compared to WT and complemented strains (>90%) in both TA and NB media (Fig. 2). The difference between TA and NB highlights a cation dependency: infection efficiency and lysis were strongly enhanced in the presence of divalent cations (TA medium), consistent with prior reports that Ca²⁺ and Mg²⁺ facilitate phage adsorption by reducing electrostatic repulsion (30,58–60). Although we did not directly measure cell surface charge, the data suggest that *gtrB*-mediated glycosylation may influence the physicochemical properties of WTA/LTA, modulating the electrostatic interactions needed for binding of phage components such as tail fibers. Comparable cation-enhanced adsorption has been described in *Listeria* and *Salmonella* phages, where divalent ions bridge negatively charged cell wall components and phage structural proteins (59,60).

Microscopy (Fig. 5A–C) revealed notable structural differences in BC2012. Gram staining suggested a Gram-negative-like phenotype (Fig. 5A), TEM showed thinner cell walls (Fig. 5B), and FE-SEM revealed smoother, less structured surfaces (Fig. 5C). While these observations clearly demonstrate altered cell morphology, the biochemical basis—potentially improper WTA/LTA glycosylation—remains to be confirmed. The smoother surfaces and apparent loss of structural features could reduce receptor density, consistent with the observed decrease in phage adsorption and larger, more diffuse plaques (Fig. 4). These phenotypic links are supported by prior studies in *S. aureus* and *Lactobacillus delbrueckii*, where glycosyltransferases modulate LTA glycosylation and D-alanylation, affecting surface charge and phage binding (61,62).

Autolysis assays indicated that BC2012 cultures underwent spontaneous lysis under static conditions but remained stable when shaken (Fig. 6A–B; Fig. 7A–B). This suggests that *gtrB* disruption compromises cell wall stability, potentially by weakening cross-links between peptidoglycan and teichoic acids. The reduced turbidity and CFU counts in static cultures (Fig. 7A) align with observations in Gram-positive mutants with defective teichoic acid synthesis, where altered glycosylation predisposes cells to stress-induced autolysis (62–67).

Taken together, our results demonstrate that disruption of *gtrB* in *B. cereus* substantially affects surface architecture, adsorption efficiency, plaque morphology, and cell wall integrity. While the precise biochemical mechanisms—such as changes in glycosylation patterns or surface charge— require further investigation, this study identifies a previously uncharacterized host determinant of phage susceptibility in *B. cereus*. Moreover, the observed cation dependency underscores the importance of electrostatic interactions in phage-host dynamics. These insights may guide the design of phage-resistant industrial strains or optimized phage therapy strategies targeting the *B. cereus* group.

## Acknowledgements

This research was supported by the Regional Innovation System & Education (RISE) initiative, funded by the Ministry of Education and administered by the National Research Foundation of Korea (NRF).

---

**Supplementary figure 1.**
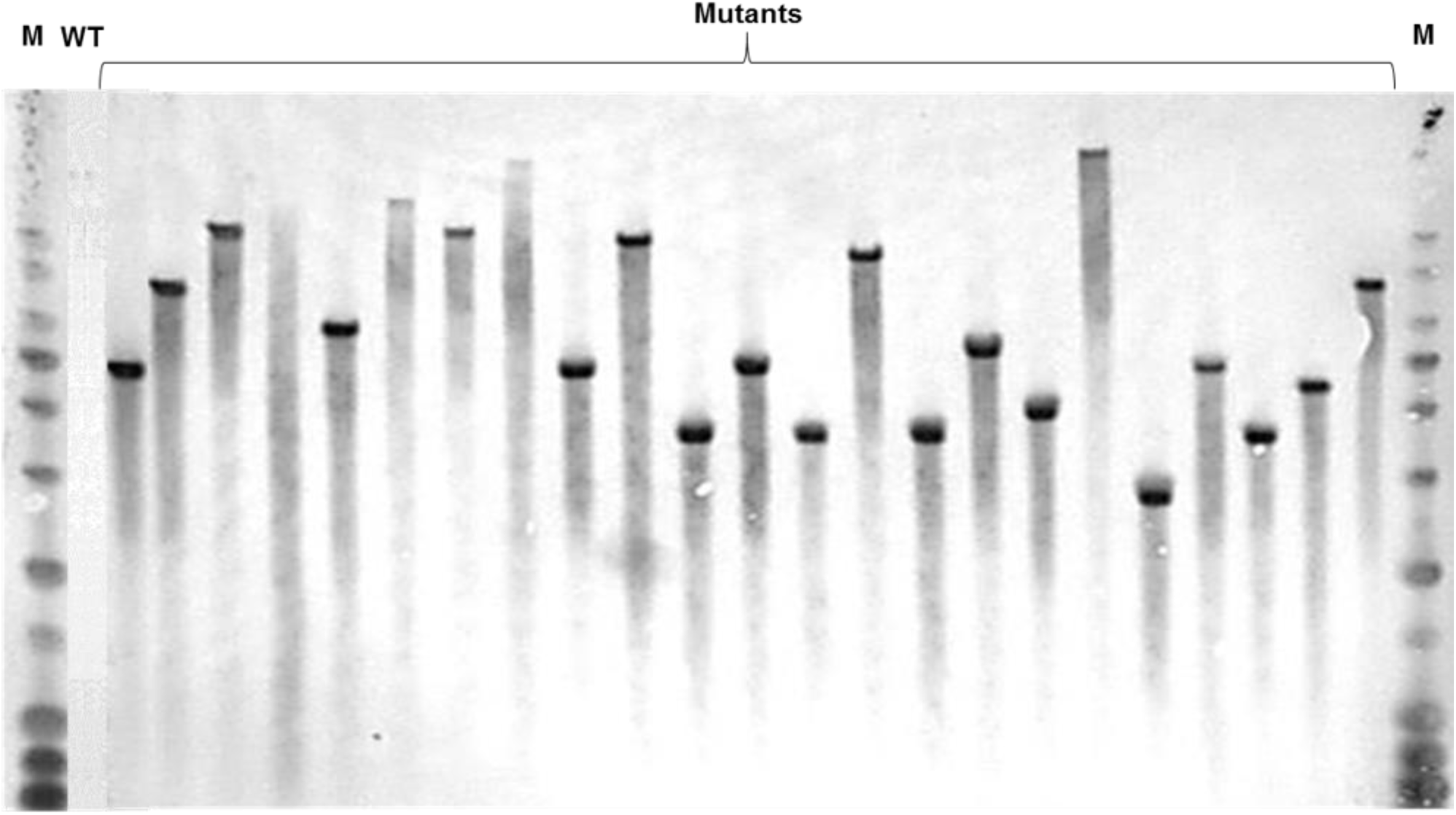
Southern hybridization using spectinomycin gene as probe. The Southern blot demonstrates randomness and single-copy insertion in 23 strains using wild-type as a control.

**Supplementary figure 2.**
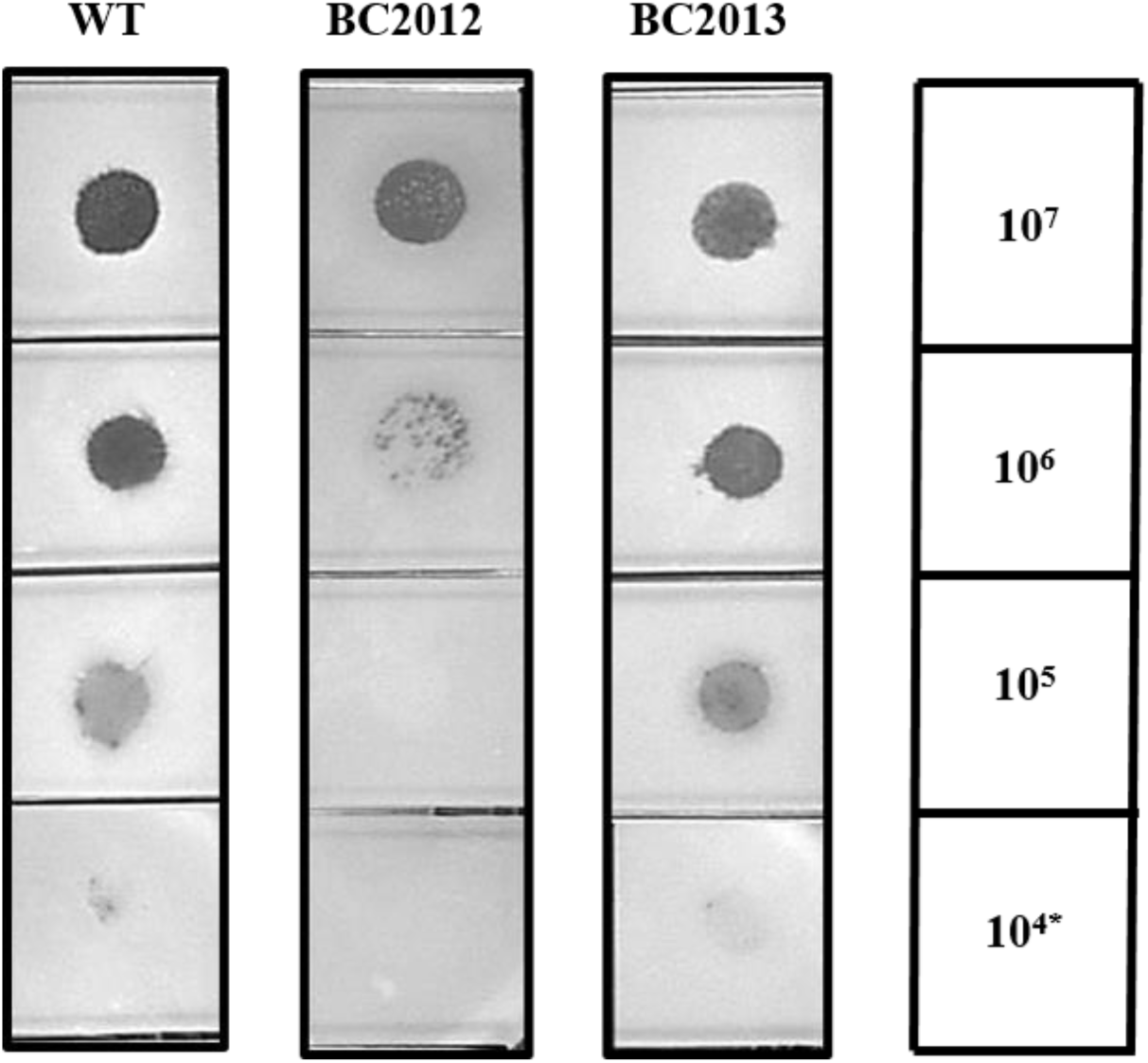
Bacterial spot assay of BCP8-2 on wild type (WT), the mutant BC2012, and the complementation strain BC2013. (5 µL of sample used)

